# Sex differences in adolescent cannabis vapor self-administration mediate enduring effects on behavioral flexibility and prefrontal microglia activation in rats

**DOI:** 10.1101/2023.01.21.524468

**Authors:** Timothy G. Freels, Sara R. Westbrook, Hayden R. Wright, Jacqulyn R. Kuyat, Erica Zamberletti, Alexandra M. Malena, Max W. Melville, Amanda M. Brown, Nicholas C. Glodosky, Darren E. Ginder, Courtney M. Klappenbach, Kristen M. Delevich, Tiziana Rubino, Ryan J. McLaughlin

## Abstract

Cannabis is the most used illicit drug in the United States. With many states passing legislation to permit its recreational use, there is concern that cannabis use among adolescents could increase dramatically in the coming years. Historically, it has been difficult to model real-world cannabis use to investigate the causal relationship between cannabis use in adolescence and behavioral and neurobiological effects in adulthood. To this end, we used a novel volitional vapor administration model to investigate long-term effects of cannabis use during adolescence on the medial prefrontal cortex (mPFC) and mPFC-dependent behaviors in male and female rats. Adolescent (35-55 day old) female rats had significantly higher rates of responding for vaporized Δ^9^-tetrahydrocannabinol (THC)-dominant cannabis extract (CAN_THC_) compared to adolescent males. In adulthood (70-110 day old), female, but not male, CAN_THC_ rats also took more trials to reach criterion and made more regressive errors in an automated attentional set-shifting task compared to vehicle rats. Similar set-shifting deficits were observed in males when they were exposed to a non-contingent CAN_THC_ vapor dosing regimen that approximated CAN_THC_ self-administration rates in females. No differences were observed in effort-based decision making in either sex. In the mPFC, female (but not male) CAN_THC_ rats displayed more reactive microglia with no significant changes in myelin basic protein expression or dendritic spine density. Together, these data reveal important sex differences in rates of cannabis vapor self-administration in adolescence that confer enduring alterations to mPFC structure and function. Importantly, female-specific deficits in behavioral flexibility appear to be driven by elevated rates of CAN_THC_ self-administration as opposed to a sex difference in the effects of CAN_THC_ vapor *per se*.

## INTRODUCTION

Cannabis is the most widely used illicit substance among adolescents and the prevalence of heavy cannabis use during adolescence has seen a threefold increase over the past 25 years. In the US, vaping of high-potency cannabis concentrates during adolescence has increased dramatically in recent years, from 24.2% in 2017 to 52.1% in 2019 among past 30-day cannabis consumers^1,2^. This is concerning because adolescence is a critical period of cortical development^3^, for which the endogenous cannabinoid system plays a prominent role^4^—thus, cannabis could interfere with these important neurodevelopmental processes, potentially increasing the risk for cognitive dysfunction later in life^5^.

Accordingly, human studies have revealed that adolescent cannabis use is associated with deficits in aspects of executive function such as impulse control, attention, visuospatial functioning and behavioral flexibility ^6–9^. For instance, adolescent-onset cannabis users show a pervasive deficit in the ability to shift strategies in the Wisconsin Card Sorting task^8^. Moreover, the earlier participants began using cannabis, the more difficulty they had learning new strategies and perseverated with the old, incorrect strategy^8^. These deficits could be related to willingness to expend effort, as both acute and chronic cannabis use have been associated with apathy, amotivation, and other reward processing deficits^10,11^. Adolescent cannabis use also has been associated with changes in cortical structure, including decreased white matter^12,13^. These data suggest that there are relationships between adolescent cannabis use and cognitive- and brain-related endpoints. However, human studies are not well suited to interrogate cause-effect relationships and as such, the long-term consequences of adolescent cannabis use remain elusive.

Preclinical studies using rodent models mirror data from human studies, demonstrating long-term cognitive impairments following injections of synthetic CB_1_R agonists^14–16^ or isolated cannabis constituents such as Δ^9^-tetrahydrocannabinol (THC)^17–22^. However, the drug and route of administration commonly used in preclinical studies raise the possibility that effects of adolescent cannabinoid exposure may not translate to human populations^23^. Whereas most preclinical studies employ forced injections of synthetic cannabinoids or high doses of isolated THC, it is well known that the most common route of administration for human users is volitional intrapulmonary intake of broad-spectrum cannabis products^24^. This is important because comparisons of injected vs. vaporized THC have revealed different pharmacokinetic profiles despite producing equal peak concentrations of circulating THC^25^. To address this translational gap, we recently developed a more ecologically relevant model of cannabis use in rats. This approach more accurately mimics the human experience of cannabis use by allowing self-administration of vaporized whole-plant cannabis extracts using commercially available e-cigarette technology^26^. Thus, this model provides a valid and reliable approach for studying the impacts of cannabis use during adolescence on executive functioning and prefrontal cortical structure in rats.

In this study, we used THC-vs. cannabidiol (CBD)-dominant vaporized cannabis extracts to test the hypothesis that self-administration of THC-dominant cannabis vapor during adolescence impairs behavioral flexibility and effort-based decision making in adulthood, with corresponding alterations in mPFC microglia activation, white matter expression, and dendritic spines.

## METHODS

### Animals

Male and female Sprague-Dawley rats obtained from Simonsen Laboratories (Gilroy, CA) on postnatal day (PND) 25 were housed in our climate-controlled vivarium on a 12:12 h reverse light cycle (lights off at 7:00) with food and water available *ad libitum*. After habituation and handling (Supplemental Material), rats underwent experimental procedures as shown in Figure 1. All experimental procedures adhered to the National Institutes of Health Guide for the Care and Use of Laboratory animals and were approved by the Washington State University Institutional Animal Care and Use Committee.

**Figure 1.**
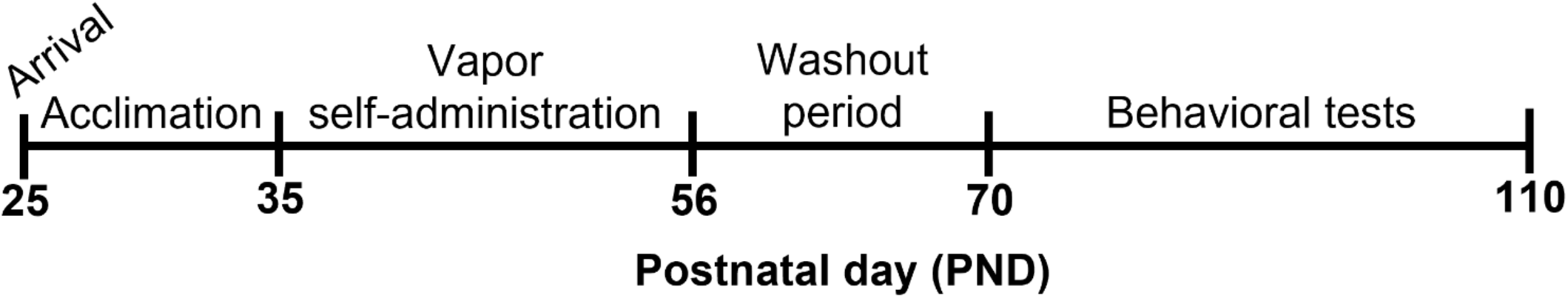
Experimental timeline. Sprague-Dawley rats of both sexes arrived at our facility on PND 25. After acclimation and handling, rats underwent daily sessions from PND 35-55 during which they could self-administer vapor from cannabis extracts with high concentrations of THC (CAN_THC_), CBD (CAN_CBD_), or vehicle (VEH). Rats were given a two-week washout period (PND 56-69) prior to tests of behavioral flexibility and effort discounting in adulthood (PND 70-110). Rats were maintained at 90% free-feeding weight during behavioral testing. Brains were collected at the end of the experiment to assess changes in the medial prefrontal cortex.

### Drugs

Natural cannabis extracts that contained high concentrations of THC (CAN_THC_) or CBD (CAN_CBD_) were acquired from the National Institute on Drug Abuse (NIDA) Drug Supply Program. The certificates of analysis provided by NIDA indicated that the CAN_THC_ extract contained 28.4% THC, 1.38% CBD, and 1.8% cannabinol, whereas the CAN_CBD_ extract contained 1.16% THC, 59.34% CBD, 2.1% cannabichromene, 1.1% cannabigerol, and <0.01% tetrahydrocannabivarin and cannabinol.

Based on previous research^26^, CAN_THC_ and CAN_CBD_ solutions were prepared at a 400 mg/ml concentration by dissolving raw cannabis extract into a vehicle (VEH) of 80% propylene glycol/20% vegetable glycerol under continuous stirring at 60°C as described previously^26^.

### Vapor Self-Administration

Cannabis vapor self-administration was conducted using second generation vapor chambers from La Jolla Alcohol Research Inc. ([LJARI] La Jolla, CA) equipped with vapor tanks (SMOK Tank TFV8 with M2 atomizers), 2 nosepoke ports, and corresponding cue lights controlled by MED Associates IV software (Fairfax, VT) as previously described^26^. Rats were trained to self-administer vapor from CAN_THC_, CAN_CBD_, or VEH preparations on a fixed ratio 1 (FR1) schedule from PND 35-55. During these 1 h sessions, a response made on the active port resulted in 3 s delivery of vapor and simultaneous illumination of a cue light located above the active port. The cue light remained illuminated for a 60 s time-out period during which active responses were recorded but had no consequence. Responses made on the inactive port were recorded but had no consequences. A discrimination index for the active port was calculated as 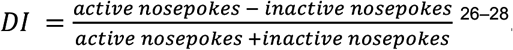. Dependent measures for vapor self-administration included nosepoke responses into the active and inactive ports, vapor deliveries, and discrimination index, and were analyzed using linear mixed models with session as the within-subject factor.

### Behavioral Testing

Beginning on PND 70, tests of attentional set-shifting and effort-based decision making were conducted in 10” x 12” x 12” (L x W x H) Coulbourn Habitest chambers (Coulbourn Instruments, Holliston, MA) using Graphic State 4 software^29,30^. Briefly, rats underwent lever shaping and retractable lever training until ≤ 5 omissions were made for 2 consecutive sessions, followed by visual cue discrimination, set-shifting, and reversal learning test phases to assess domains of behavioral flexibility as described previously^33,34^. Next, rats were retrained on retractable lever training and effort discounting was assessed^30^. See Supplemental Material for a complete description of behavioral testing procedures.

### Immunofluorescence for Medial Prefrontal Cortex Myelination and Microglia Morphology

Brains were collected following transcardial perfusion, post-fixed in 4% paraformaldehyde overnight, sunk in 30% sucrose solution, and stored in 1X PBS + sodium azide until needed. Brains from a subset of animals (n=3/group) were stained for MBP and Iba1 (Supplemental Material) to assess myelination and microglia morphology, respectively. For MBP analysis, mean grey value from four different slices containing the mPFC for each animal was quantified using ImageJ software (NIH, Bethesda, MD, United States). For Iba1 staining of microglia, four parameters were analyzed as described previously^31^: cell surface area, cell perimeter, soma area, and roundness of the soma. Since microglia are known to mediate synaptic pruning, another subset of rats was used to assess effects of vapor self-administration during adolescence on dendritic spines in the mPFC (Supplemental Material).

### Statistical Analyses

Statistical outliers were determined via the ROUT method (Q = 1%) in GraphPad Prism software (version 9.4.1, San Diego, CA, USA) and removed prior to statistical analyses (Supplemental Material). The alpha level was set to *p* < 0.05, and dependent measures (Supplemental Material) were assessed using analyses of variance (ANOVAs) and linear mixed models (where appropriate) separately for each sex with the between-subjects factor of treatment. Significant main effects of treatment were further examined using Dunnett’s post-hoc tests with vehicle as the control. All other significant main effects or interactions were followed up with Tukey post-hoc tests. Effect size estimates were calculated using Cohen’s *d*. All statistical analyses were conducted in SAS (version 9.4, SAS Institute Inc., Cary, NC, USA) using PROC GLIMMIX.

## RESULTS

### Adolescent female rats self-administer more THC- (but not CBD-) dominant cannabis vapor than male rats

Adolescent rats of both sexes earned a similar number of CAN_THC_ and vehicle vapor deliveries [Treatment by Sex: F_2,82_ = 10.24, *p* = 0.0001, Figure 2A,B], and earned more vehicle than CAN_CBD_ (females: *p* < 0.0001, *d* = 1.33; males: *p* = 0.0003, *d* = 0.77). Females earned more vehicle (*p* < 0.0001, *d* = 0.68) and CAN_THC_ (*p* < 0.0001, *d* = 0.95) vapor deliveries than their male counterparts. Sex differences were also seen in active responses, inactive responses, and discrimination indices (Supplemental Material, Figure S1). Given the significant effect of sex on all self-administration measures, data were stratified by sex for all subsequent statistical analyses.

**Figure 2.**
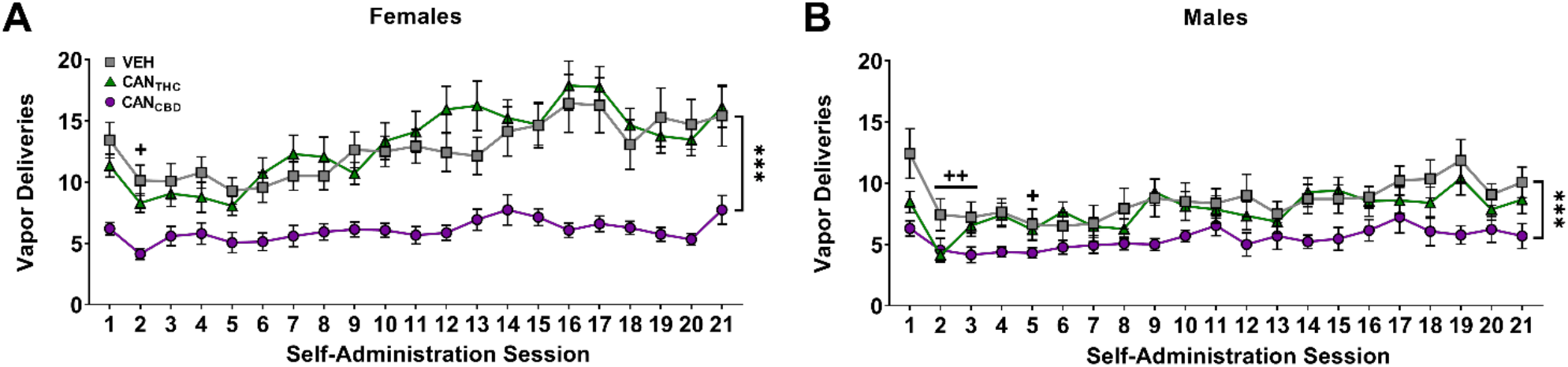
Adolescent rats of both sexes self-administer vapor from cannabis extracts with high concentrations of THC (CAN_THC_), CBD (CAN_CBD_), or vehicle (VEH). Females (**A**) self-administered significantly more CAN_THC_ and vehicle vapor compared to males (**B**) and thus all further analyses were stratified by sex. Data are shown as mean (± SEM). n = 13-17/group. ^++^*p* < 0.01 vs. session 1, collapsed across treatment and sex; \*\*\**p* < 0.001 vs. VEH, collapsed across session and sex.

### Adolescent cannabis self-administration impairs behavioral flexibility in female rats in adulthood

After a two-week washout period, behavioral flexibility was assessed in visual cue discrimination (set), set-shifting, and reversal tasks. For females, treatment did not significantly impact trials to reach criterion during the initial visual discrimination (Figure 3A) or the reversal tasks (Figure 3C). However, during the set-shift (Figure 3B), females who self-administered CAN_THC_ vapor during adolescence took significantly more trials to reach criterion compared to vehicle females (*p* = 0.0065, *d* = 1.63), [Treatment: F_2,35_ = 4.91, *p* = 0.013]. This increase in trials to criterion was driven by the greater number of regressive errors made by CAN_THC_ females compared to vehicle females (*p* = 0.0057, *d* = 1.68), [Treatment by Error Type interaction: F_4,38_ = 3.41, *p* = 0.018, Figure 3D]. Female CAN_THC_ rats made significantly more regressive errors relative to the other two error types—never-reinforced (*p* =0.0017, *d* = 0.92) and perseverative (*p* = 0.0067, *d* = 1.26). For males, there were no significant effects of treatment on trials to reach criterion for the initial visual cue discrimination, the set-shift, or the reversal. During the set-shift, males, regardless of treatment, made significantly more regressive errors than never-reinforced errors (*p* = 0.012, *d* = 0.50), [Error Type: F_2,36_ = 9.08, *p* = 0.0006]. Omissions during each of the tasks were also analyzed (Supplemental Material, Figure S2).

**Figure 3.**
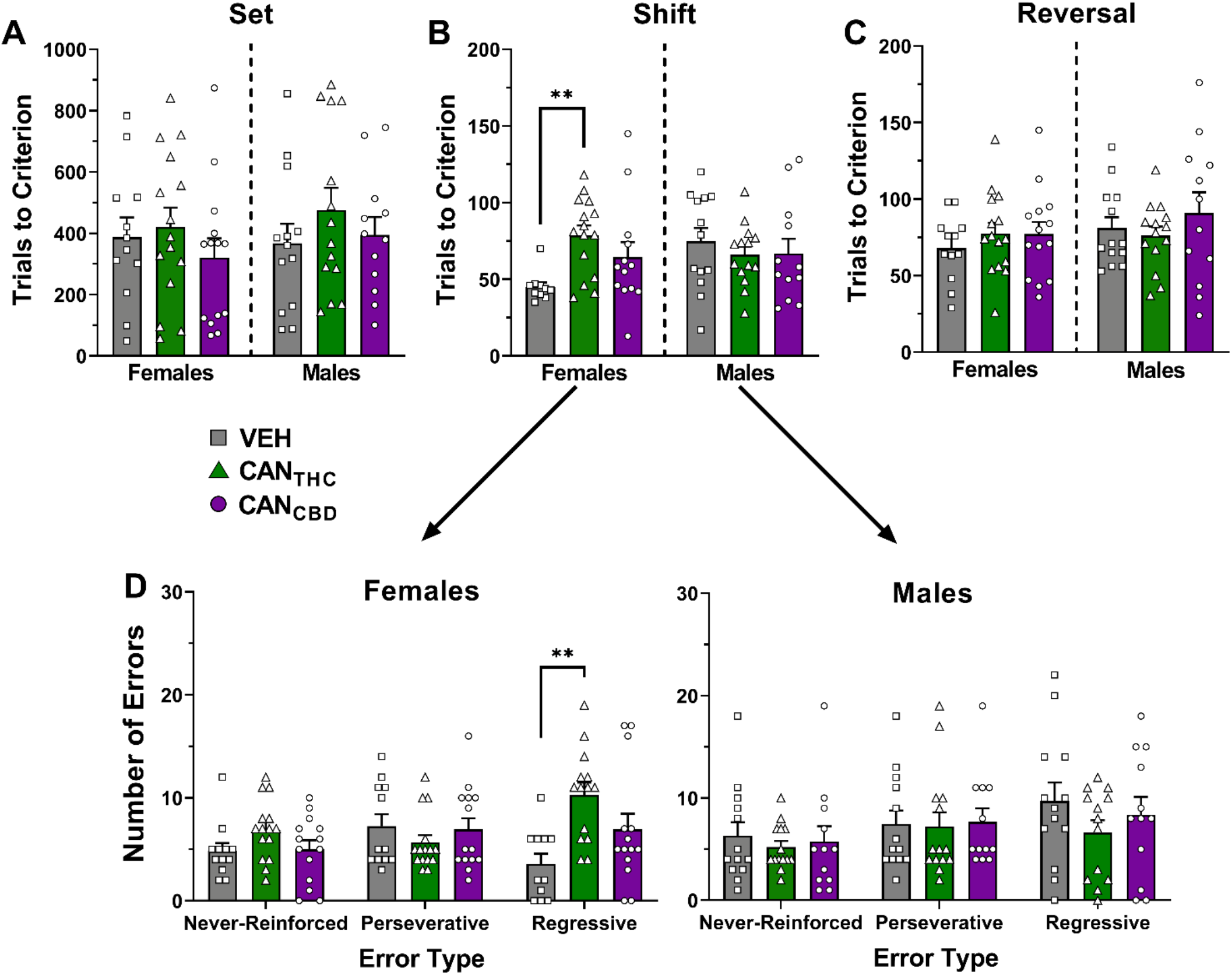
CAN_THC_ vapor during adolescence impairs behavioral flexibility in female rats when tested in adulthood. Trials to reach criterion in the visual cue discrimination (set; **A**), set-shifting (**B**), and reversal (**C**) tasks in rats of both sexes. (**D**) Number of different types of errors made during the set-shifting task. Data are shown as mean (± SEM). n = 10-15/group. \*\**p* < 0.01 vs. VEH.

Since females earned more CAN_THC_ vapor than males, behavioral flexibility was also assessed in a separate cohort of adult rats that were exposed during adolescence to non-contingent CAN_THC_ or vehicle vapor at quantities comparable to that self-administered by adolescent females (Supplemental Material, Figure S3). Non-contingent CAN_THC_ vapor exposure produced behavioral flexibility deficits in males similar to those seen after self-administration in females. In contrast, there were no significant effects of vapor self-administration on effort discounting for either sex (Supplemental Material, Figure S4).

### Adolescent cannabis vapor self-administration produces sex-dependent effects on microglia morphology (but not myelination) in adulthood

*Microglia*. Morphological changes in microglia in the mPFC were visualized with Iba1 fluorescence to determine if microglia are altered in adulthood following vapor self-administration during adolescence (Figure 4A). In females, there were significant effects of treatment on all four measures— cell surface area [F_2,6_ = 100.31, *p* < 0.0001, Figure 4B], cell perimeter [F_2,6_ = 191.17, *p* < 0.0001, Figure 4C], soma area [F_2,6_ = 18.05, *p* = 0.0029, Figure 4D], and soma roundness [F_2,6_ = 14.91, *p* = 0.0047, Figure 4E]. Female CAN_THC_ rats had significantly reduced microglia surface area (*p* < 0.0001, *d* = 10.28), reduced microglia perimeter (*p* < 0.0001, *d* = 14.03), larger soma area (*p* = 0.0019, *d* = 4.54), and rounder soma (*p* = 0.0055, *d* = 3.35) compared to vehicle females. Female CAN_CBD_ rats had significantly reduced cell perimeter (*p* = 0.005, *d* = 3.28) but no other significant microglial changes relative to vehicle females.

**Figure 4.**
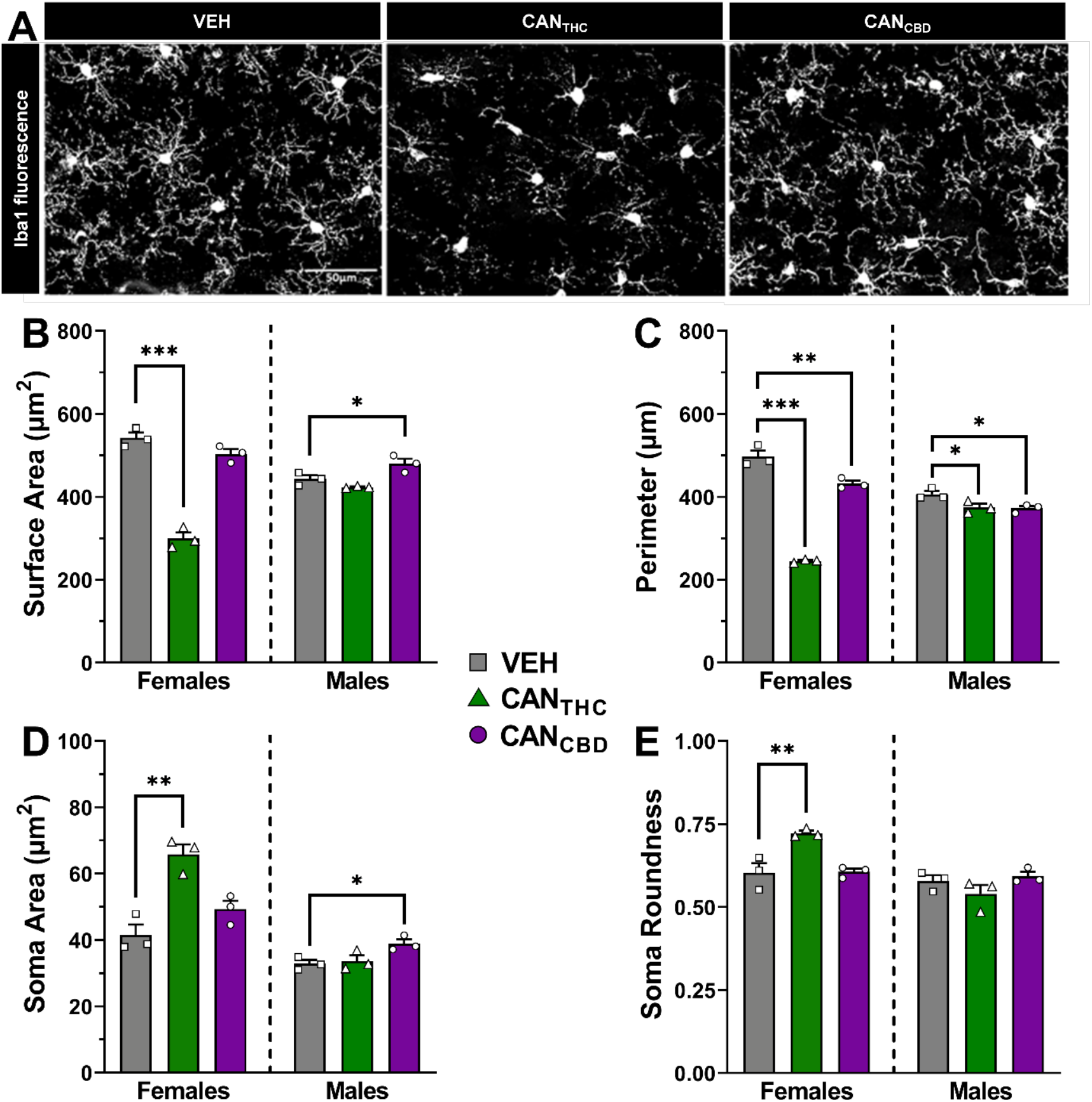
Adolescent cannabis vapor self-administration affects mPFC microglial morphology in adulthood in a sex- and cannabis vapor-specific manner. In females, CAN_THC_ vapor self-administration during adolescence leads to more reactive microglia in the mPFC in adulthood. In males, microglia morphological changes in the mPFC occur following CAN_CBD_ vapor self-administration in adolescence. (**A**) Representative images of microglia stained for ionized calcium binding adaptor molecule 1 (Iba1) in the mPFC. Microglia cell (**B**) surface area and (**C**) perimeter, as well as soma (**D**) size and (**E**) roundness in adult rats of both sexes with a history of vapor self-administration during adolescence. Data are shown as mean (± SEM). n = 3/group. \**p* < 0.05, \*\**p* < 0.01, \*\*\**p* < 0.001 vs. VEH.

Males also exhibited significant effects of treatment on microglial morphology, including cell surface area [F_2,6_ = 10.99, *p* = 0.0099, Figure 4B], cell perimeter [F_2,6_ = 6.64, *p* = 0.0302, Figure 4C], and soma area [F_2,6_ = 5.60, *p* = 0.0425, Figure 4D]; however, there were no significant effects on soma roundness (Figure 4E). In contrast to females, the significant changes seen in males were primarily in CAN_CBD_ rats relative to vehicle vapor. Male CAN_CBD_ rats had significantly increased microglial surface area (*p* = 0.0410, *d* = 2.02), reduced cell perimeter (*p* = 0.0302, *d* = 2.85), and increased soma area (*p* = 0.0374, *d* = 2.87) compared to vehicle males. Cell perimeter was also significantly reduced in CAN_THC_ males relative to vehicle males (*p* = 0.0403, *d* = 2.28). Since microglia are known to mediate synaptic pruning, we also examined the effects of vapor self-administration during adolescence on dendritic spines in the mPFC in adulthood, but there were no significant effects (Supplemental Material, Figure S5).

#### Myelination

MBP fluorescence was measured to assess effects of vapor self-administration on myelination in the mPFC (Figure 5A). For both males and females, there were no significant effects of treatment on MBP Grey Value (Figure 5B).

**Figure 5.**
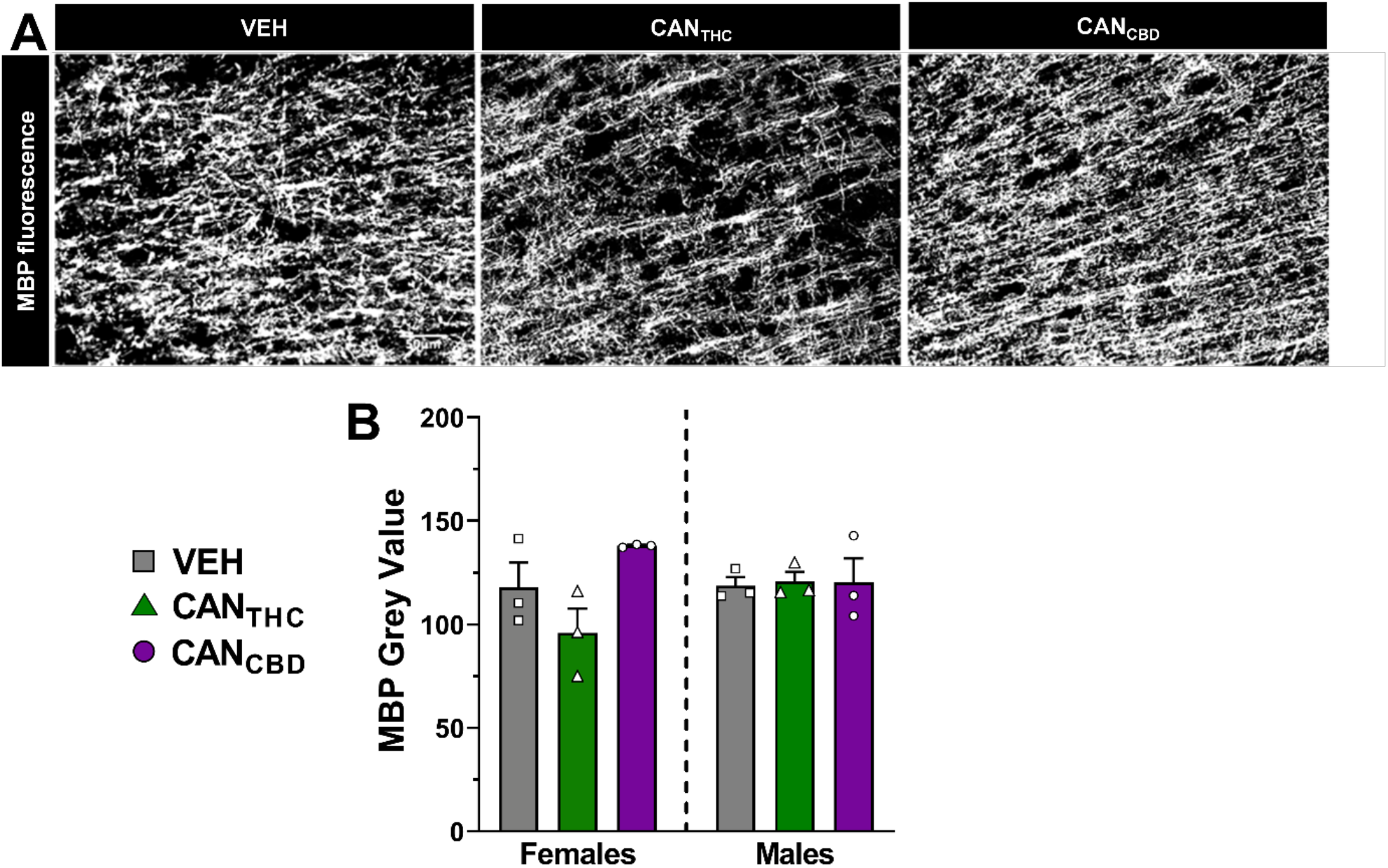
Myelination in the mPFC in adulthood is not significantly altered by cannabis vapor self-administration during adolescence. (**A**) Representative images of myelin stained for myelin basic protein (MBP) in the mPFC. (**B**) MBP grey value for adult rats of both sexes with a history of vapor self-administration during adolescence. Data are shown as mean (± SEM). n = 3/group.

## DISCUSSION

In the present study, male and female rats self-administered vehicle or cannabis vapor dominant in THC or CBD during adolescence with behavioral and biological endpoints assessed in adulthood following a two-week washout period. Adolescent female rats earned more CAN_THC_ and vehicle vapor deliveries than adolescent males. Adolescent rats self-administered both CAN_THC_ and vehicle vapor more readily than CAN_CBD_. Since cannabis use in humans has been associated with executive dysfunction and amotivation, we used an attentional set-shifting task^32^ and an effort discounting task^33^ to assess behavioral flexibility and amotivation, respectively. Our findings partially supported our hypothesis, as females, but not males, that self-administered CAN_THC_ vapor committed more regressive errors leading to impaired shifting between strategies, while effort-based decision making was not altered in rats of either sex.

Our finding that adolescent females self-administer more CAN_THC_ vapor than adolescent males is consistent with our previous report of sex differences in adult rats^28^, suggesting that sex differences in vapor self-administration are present as early as adolescence and persist into adulthood. Interestingly, adolescent rats self-administer CAN_CBD_ to a lower extent than CAN_THC_ vapor, consistent with our prior work in adult males^26^, suggesting that the phytocannabinoid concentrations present in the cannabis extract influence vapor self-administration. It is important to note that both adolescent males and females readily self-administer the vehicle vapor to similar levels as the CAN_THC_ vapor. This finding is in line with our previous report in adult rats of both sexes, where differences in responding for cannabis vapor vs. vehicle vapor only emerged after increasing the demand of the reinforcement schedule^28^. Notably, while CAN_THC_ and vehicle vapor were self-administered at similar levels during adolescence, only CAN_THC_ vapor self-administration induced behavioral deficits in adulthood in female rats.

After vapor self-administration in adolescence, deficits in behavioral flexibility, but not effort-based decision making, were observed in adulthood. Specifically, CAN_THC_ females were impaired when shifting strategies compared to vehicle females. This deficit resulted from increased regressive errors made during the shift, indicating that CAN_THC_ females were impaired in maintaining the new strategy. When we passively exposed adolescent males to the same quantity of CAN_THC_ vapor that adolescent females self-administered, we found the same deficits in the shift component in males. This finding suggests that the *quantity* of CAN_THC_ exposure during adolescence is a primary driver of CAN_THC_-induced behavioral flexibility deficits, rather than sex differences in the effects of CAN_THC_ vapor. Accordingly, a recent study found that adolescent male rats needed a higher dose of injected THC (5 mg/kg vs. 1.5 mg/kg) to produce executive dysfunction when tested in adulthood^34^. Notably, we did not observe behavioral flexibility deficits in passively exposed female rats (see Figure S3). Though unexpected, these data could suggest that the unintended stress of forced cannabis vapor delivery have sex-differentiated effects that exacerbate cannabis-induced impairments in adolescent male rats but attenuate those impairments in female rats. In line with this possibility, pubertal stress has been shown to induce anhedonia in adult males but not females, which is paralleled by a male-specific decrease in striatal dopamine turnover ^35^.

Another aim was to test whether cannabis vapor self-administration during adolescence disrupts ongoing development of the mPFC. During adolescence, the brain undergoes extensive development and reorganization of cortical regions that are critical for executive functioning^3,36,37^. These developmental processes include myelination^38^ and synaptic pruning^39^, which is partly mediated by microglia^40,41^. Supporting the endocannabinoid (eCB) system’s vital role in mPFC development, injections of THC during adolescence have been shown to disrupt these processes^21,42–44^. Interestingly, our observed behavioral deficits following CAN_THC_ vapor in adolescence were specific to the shift component of the behavioral flexibility task, which has been shown to be mPFC-dependent^32^. We predicted that adolescent use of cannabis vapor would decrease myelination but increase microglial activation and synaptic pruning, thereby producing a net reduction in dendritic spine density. While we found no effects of vapor self-administration on myelination, we did find that mPFC microglia of female CAN_THC_ rats were more amoeboid-like—characteristic of the reactive state. This finding is consistent with previous reports in females using THC injections^21,42^. Interestingly, Zamberletti et al.^21^ found that preventing the altered microglia phenotype through co-administration of ibudilast (a compound that blocks microglia activation) and THC rescued THC-induced behavioral deficits in adulthood, suggesting that more reactive microglia may play a causal role in THC-induced behavioral dysfunction in adolescent females. Future work should determine whether more reactive microglia also contribute to the behavioral flexibility deficits we observed.

More reactive microglia may lead to premature and/or greater synaptic pruning during adolescence, potentially resulting in fewer dendritic spines in the mPFC. In our analysis of dendritic spines, we did not observe a significant decrease in spine density in our CAN_THC_ group. This is consistent with a recent study in male rats that reported premature synaptic pruning (but not greater pruning overall) immediately following adolescent THC exposure that did not manifest as a difference in spine number after a washout period^44^. Premature synaptic pruning may lead to aberrant synaptic connections, which could then influence cognitive function in adulthood without a change in the total number of spines. However, it is important to note that we observed more reactive microglia only in females, and our spine analyses were not powered to detect sex differences due to issues with cell filling in brain slices taken from male rats. Thus, further investigation is needed to determine if the more reactive microglia seen in females are associated with premature and/or greater synaptic pruning.

In conclusion, the current study revealed sex differences in cannabis vapor self-administration in adolescence that produced behavioral inflexibility and alterations in microglia morphology later in life, which are in line with effects that have been observed following injections of low doses of THC. However, by using a more translational model that employs response-contingent vapor delivery, we uncovered the importance of the *quantity* of THC exposure and volitional control over vapor delivery during adolescence as primary exacerbating factors that influence long-term behavioral and biological outcomes.

## ACKNOWLEDGEMENTS

The authors would like to thank Maury Cole and LJARI, Inc. for their continued support with vapor chamber troubleshooting and optimization. The authors would also like to thank Abigail Rossi, Alyssa Hampton, and Manuel Rojas for their assistance with behavioral testing and tissue analysis.

## AUTHORSHIP CONTRIBUTION STATEMENT

**TGF:** data curation (equal), investigation (lead), project administration (lead), software (equal); **SRW:** data curation (equal), formal analysis (lead), writing-original draft (lead), writing-review & editing (equal); **HRW:** investigation (supporting), project administration (supporting), software (equal), writing-review & editing (supporting); **EZ:** formal analysis (supporting), investigation (supporting), visualization (supporting); **JRK:** investigation (supporting), project administration (supporting), writing-review & editing (supporting); **ANM:** investigation (supporting); **MWM:** investigation (supporting); **AMB:** investigation (supporting); **NCG:** investigation (supporting); **DEG:** investigation (supporting); **CMK:** formal analysis (supporting), visualization (supporting); **KMD:** resources (equal), software (equal), supervision (supporting), writing-review & editing (supporting); **TR:** conceptualization (supporting), resources (equal), software (equal), supervision (supporting); **RJM:** conceptualization (lead), funding acquisition (lead), methodology (lead), resources (lead), supervision (lead), visualization (supporting), writing-original draft (supporting), writing-review & editing (equal)

## CONFLICT OF INTEREST STATEMENT

The authors have no competing personal or financial interests or any other conflicts of interest to disclose.

## FUNDING STATEMENT

This study was supported by an NIH R21 grant from the National Institute on Drug Abuse (R21DA043722-01) awarded to RJM, as well as funds provided for medical and biological research by the State of Washington Initiative Measure No. 171 (RJM). EZ and TR are supported by funds from the Zardi-Gori Foundation (zamb001zardigori).

## Notes

### Competing Interest Statement

The authors have declared no competing interest.

