## Supplemental Files for "Sex differences in adolescent cannabis vapor self-administration mediate enduring effects on behavioral flexibility and prefrontal microglia activation in rats"

**SUPPLEMENTAL MATERIAL**

### METHODS

#### Acclimation

Rats arrived on postnatal day (PND) 25 and were given 1 week to habituate to our housing conditions. Following habituation, rats were handled for a minimum of 3 min for 3 consecutive days before testing to acclimate them to experimenters.

#### Attentional Set Shifting

*Pretraining*. Pretraining consisted of lever shaping, retractable lever training, and side preference testing. During lever shaping, rats were trained to press either the left or right lever (counterbalanced across rats) on a FR1 reinforcement schedule. Rats were required to make ≥ 60 presses on a given lever during a 30 min session. Once criterion was reached, rats were shaped on the opposite lever before advancing to retractable lever training. In retractable lever training, the left or right lever was psudorandomly extended every 20 s, and rats were required to press the extended lever within 10 s to receive a reward. If no response occurred within 10 s, levers were retracted, and the trial was scored as an omission. Rats performed retractable lever training for a minimum of 5 days. Training criterion was met when rats made ≤ 5 omissions for 2 consecutive sessions. Side preference testing was conducted on the same day rats completed retractable lever training. Side preference testing consisted of 7 trial blocks (2-8 subtrials) with a 20 s intertrial interval. For each initial trial of the 7 trial blocks, rats had to select the left or right lever to receive a reward. During subtrials, only responses on the lever opposite of the initial choice were reinforced. The lever that the rats selected for the majority of initial responses across the 7 trial blocks was considered the side preference.

*Testing Phases.* Next, rats advanced to the visual cue discrimination, set shifting, and reversal testing phases. Criterion for completion of each task was 10 correct consecutive responses during a session. In the visual cue discrimination task, rats were required to always press the lever indicated by an illuminated cue light to receive a reward. During the set-shifting task, an extradimensional shift was introduced, such that rats had to disregard the previous cue-based rule and only press the lever opposite of their side preference. For the reversal task, rats were required to make an intradimensional shift by always pressing the lever opposite of the lever that was reinforced during the set-shifting task to receive a reward. Assessment of behavioral flexibility was based on the total number of trials rats needed to reach criterion during visual cue discrimination, set shifting, and reversal tasks. Trials to criterion was analyzed using separate one-way (treatment) ANOVAs for each task—set, shift, and reversal.

*Errors and Omissions*. Error types were tabulated for set-shifting and reversal tasks in blocks of 16 trials. In the set-shifting task, errors were scored as “perseverative” until fewer than 6 errors were made in a trial block. Subsequent errors were then scored as “regressive”. “Never-reinforced” errors were tabulated when a rat made a response that was not reinforced in visual cue discrimination or set-shifting tasks (i.e., selection of the side preference lever when no cue light was lit above it). For the reversal task, errors were scored as perseverative until fewer than 10 were committed within a trial block, after which errors were scored as regressive. A linear mixed model with error type (Never-reinforced, Perseverative, or Regressive) as the within-subjects factor was conducted on the errors during the set-shifting task. Additionally, omissions were tabulated during each task, serving as an index of gross motivation and were analyzed using separate one-way (treatment) ANOVAs for each task.

**Effort Discounting**

The effort discounting task was adapted from Simon et al. (2013). All rats performed the effort discounting task following completion of the behavioral flexibility task. In the effort discounting task, one lever was assigned as the low reward/low effort lever and the other as the high reward/high effort lever. During each 32 min session, rats completed 5 blocks of 12 trials for a total of 60 trials. Trials occurred every 40s and began with the illumination of the house light followed by the extension of one lever (forced-choice trial) or both levers (free-choice trial). Each trial block began with 2 forced-choice trials followed by 10 free-choice trials. During forced-choice trials, the low or high reward lever was presented in a random order. These trials served to establish reward receipt contingencies for each block. For free-choice trials, a single press on the low reward lever resulted in the retraction of both levers and the delivery of 1 food pellet. If the high reward lever was selected, the low reward lever was retracted and the high reward lever remained extended until the requisite number of lever presses was completed, resulting in the retraction of the high reward lever and delivery of 4 food pellets. The high reward lever was also retracted if a rat did not complete the required number of lever presses in 25 s, but the choice of the high reward lever was still recorded. The number of required presses to obtain the high reward was increased across trial blocks (i.e., 1, 2, 5, 10, and 20 presses). If no initial response was made within 25 s, the trial was scored as an omission. Rats were trained in the effort discounting task until, as a group, they exhibited selection of the high reward lever on at least 70% of trials during the first test block. Additionally, rats had to exhibit stable patterns of effort discounting across 3 consecutive sessions such that a one-way ANOVA revealed no main effect of session or session x test block interaction on high reward choice with $p$'s > 0.1 (Ghods-Sharifi & Floresco, 2010). Effort discounting was evaluated based on the preference of the rats for the high-effort/high-reward option (i.e., % high reward lever choices) during free-choice trials across test blocks and was analyzed with a linear mixed model including the within-subject factor of response requirement (1, 2, 5, 10, or 20).

**Immunofluorescence for mPFC Myelination and Microglia Morphology**

Free floating sections (40 μm thick) containing the mPFC were washed three times in 0.05% Tryton X-100 in PBS, incubated with 3% normal goat serum, 0.05% Tryton X-100 in PBS for 45 min at room temperature and then overnight at 4°C with rabbit anti-IBA1 antibody (1:1000, Wako, Neuss, Germany) diluted in blocking solution. On the second day, sections were washed and incubated for 4h at room temperature with Alexa Fluor 594 goat anti-rabbit antibody (1:2000; Invitrogen, Eugene, OR, USA) and then incubated with mouse monoclonal anti-myelin basic protein (MBP) antibody (1:500;

Immunological Sciences, Rome, Italy) in blocking solution overnight at 4 °C. After washing, signal was revealed by incubating sections with Alexa Fluor 488 goat anti-mouse antibody (1:2000; Invitrogen, Eugene, OR, USA) for 4h at room temperature. After several washes in PBS, sections were mounted onto Superfrost slides, dehydrated and coverslipped. For each animal, a complete series of one-in-six sections (240 mm apart) through the PFC was analyzed.

Digital Images were captured using Retiga R1 CCD camera (QImaging, Surrey, BC, Canada) attached to an Olympus BX51 (Tokyo, Japan) polarizing/light microscope. Ocular imaging software (QImaging, Surrey, BC, Canada) was used to import images from the camera. Images of microglia cells in the mPFC were acquired by first delineating the brain sections and the region of interest at low magnification (4x objective) and the region of interest outlines were further refined under a 40x objective for microglia analysis or under a 20x objective for analysis of MBP staining. Three rats per each experimental group (four sections/rat) were analyzed. Microglia cells were selected and cropped according to the following criteria: (1) random selection in the mPFC; (2) no overlapping with neighboring cells; (3) complete soma and branches. Selection was done blinded to the treatment. A total of twelve cells were analyzed from each animal. Each grayscale single cell cropped image was processed in a systematic way to obtain binary image using the same threshold for all pictures. The binary image was edited to clear the background and transformed into a filled shape and its pairwise outline shape that were used for morphological parameters measurements. Analysis was performed using FIJI free software (NIH, Bethesda, MD, United States). A one-way (treatment) ANOVA was conducted on the mean grey value for MBP fluorescence in the PFC. For the measures of PFC microglia morphology—cell surface area, cell perimeter, soma size, and soma roundness—one-way (treatment) ANOVAs were performed.

**Dendritic Spines in the Medial Prefrontal Cortex**

Brains from a subset of animals (n=4/group) from the self-administration study were used to examine dendritic spines of pyramidal neurons in layer II/III of the mPFC. Cells were filled using Lucifer

Yellow dye as previously described (Anderson et al., 2016; Karatsoreos et al., 2011) to visualize dendrites and spines. Dendrites in the mPFC were traced and spines were scored in Neurolucida 360. Spine type characterization was achieved via Neurolucida's default classifications of filopodia, mushroom, stubby, and thin based on published methods (Rodriguez et al., 2008). Dependent measures for dendritic spines in the PFC included spine density, spine volume, and spine length, and one-way (treatment) ANOVAs were conducted on these measures. Proportion of the different types of spines (filopodia, mushroom, stubby, and thin) was analyzed using a linear mixed model with spine type as the within-subjects factor.

**Passive Vapor Exposure**

A separate cohort of adolescent male (n=32) and female (n=32) Sprague-Dawley rats received 21 sessions of non-contingent vehicle or $CAN_{THC}$ vapor deliveries equal to the mean vehicle or $CAN_{THC}$ vapor deliveries earned by the adolescent females during each self-administration session as shown in Figure 1E. The daily sessions occurred from P35-P55. After a washout period, rats received the same behavioral flexibility assessment as described in the main text from P70-P110. Number of trials to reach criterion was analyzed separately for each sex using one-way (treatment) ANOVAs for each task—set, shift, and reversal. A linear mixed model with error type (Never-reinforced, Perseverative, or Regressive) as the within-subjects factor was conducted on the errors during the set-shifting task.

### RESULTS

**Vapor Self-administration**

Adolescent female rats made more active pokes for $CAN_{THC}$ ($p < 0.0001$, $d = 0.66$) and vehicle vapor than males ($p < 0.0001$, $d = 0.48$). However, both sexes poked significantly more for vehicle compared to $CAN_{THC}$ (females: $p = 0.0072$, $d = 0.30$; males: $p = 0.023$, $d = 0.56$) or $CAN_{CBD}$ (females: $p < 0.0001$, $d = 0.99$; males: $p = 0.0009$, $d = 0.83$), [Treatment by Sex: $F_{2,82} = 5.70$, $p = 0.0048$,

Supplemental Figure S1A,B]. Adolescent rats, regardless of sex, made significantly more nose poke responses in the first session compared to subsequent sessions with more pokes in the active port during session 1 compared to sessions 2-7 ($p$'s < 0.05, range of $d$'s = 0.58 – 0.80) [Session: $F_{20,1640}$ = 3.56, $p$ < 0.0001], and more pokes in the inactive port during session 1 compared to sessions 3-21 ($p$'s < 0.05, range of $d$'s = 0.35 – 0.86), except session 17 and 20 [Session: $F_{20,1640}$ = 2.67, $p$ < 0.0001, Supplemental Figure S1C,D]. Vehicle females made more inactive pokes than vehicle males ($p$ = 0.011, $d$ = 0.41) [Treatment by Sex: $F_{2,82}$ = 3.38, $p$ = 0.039], and vehicle males made more inactive pokes than $CAN_{THC}$ males ($p$ = 0.021, $d$ = 0.43). Adolescent rats poked more in the active port than inactive port (discrimination index > 0) after just one week of vapor self-administration (session 1 vs. session 8-21: $p$'s < 0.01, range of $d$'s = 0.49 – 0.82), [Session: $F_{20,1640}$ = 17.15, $p$ < 0.0001]. Females exhibited higher discrimination indices than males [Sex: $F_{1,82}$ = 6.64, $p$ = 0.012, $d$ = 0.39], but treatment did not influence discrimination index (Supplemental Figure S1E,F).

**Behavioral Flexibility**

*Outliers*. ROUT outlier analyses identified 2 outliers for trials to criterion during the set shift (Female VEH: n=2) and 3 outliers in regressive errors (Female VEH: n=1, Female $CAN_{THC}$: n=1, Male $CAN_{THC}$: n=1). For omissions during behavioral flexibility, 7 outliers were identified for the visual cue discrimination (Female VEH: n=1, $CAN_{CBD}$: n=2; Male VEH: n=1, $CAN_{THC}$: n=2, $CAN_{CBD}$: n=1), 10 outliers for the set-shift (Female VEH: n=2, $CAN_{THC}$: n=2, $CAN_{CBD}$: n=2; Male VEH: n=2, $CAN_{THC}$: n=1, $CAN_{CBD}$: n=1), and 8 outliers for the reversal tasks (Female VEH: n=2, Female $CAN_{THC}$: n=3; Male $CAN_{THC}$: n=1, $CAN_{CBD}$: n=2). In the rats exposed to non-contingent vapor during adolescence, ROUT outlier analyses on trials to criterion during behavioral flexibility identified 1 outlier for the visual cue discrimination (Female VEH: n=1), 3 outliers for the set-shift (Female VEH: n=1; Male VEH: n=1, $CAN_{THC}$: n=1), and 4 outliers for the errors during the set-shift (Female VEH Never-Reinforced: n=1, Female VEH Perseverative: n=1; Male VEH Regressive: n=1, Male $CAN_{THC}$ Regressive: n=1). These data points were removed, and statistical analyses conducted on the remaining data.

*Omissions*. During the visual cue discrimination task (Supplemental Figure S2A), females who self-administered $CAN_{THC}$ vapor during adolescence had significantly more omissions than their vehicle vapor counterparts ($p$ = 0.02, $d$ = 0.92), [Treatment: $F_{2,35}$ = 4.79, $p$ = 0.015]. There were no effects of treatment in males during visual cue discrimination. For rats of both sexes, there were significant effects of treatment on omissions during the set-shifting task [Females: $F_{2,32}$ = 3.50, $p$ = 0.042; Males: $F_{2,32}$ = 9.14, $p$ = 0.0007, Supplemental Figure S2B]. Post-hoc tests were not significant in females, but males who self-administered $CAN_{CBD}$ vapor during adolescence made significantly more omissions than males that self-administered vehicle vapor ($p$ = 0.0018, $d$ = 1.23). During the reversal task (Supplemental Figure S2C), adult rats of both sexes with a history of $CAN_{CBD}$ vapor self-administration during adolescence committed more omissions than their vehicle vapor counterparts (females: $p$ = 0.028, $d$ = 0.86; males: $p$ = 0.026, $d$ = 0.87), [Females: $F_{2,33}$ = 4.78, $p$ = 0.015; Males: $F_{2,33}$ = 4.33, $p$ = 0.021].

*Following Non-Contingent Vapor Exposure*. In males, treatment history did not significantly affect trials to reach criterion during visual cue discrimination (Supplemental Figure S3A) or during the reversal (Supplemental Figure S3C) tasks. During the set-shift (Supplemental Figure S3B), males with a history of passive exposure to $CAN_{THC}$ vapor took more trials to reach criterion than those exposed to vehicle vapor [$F_{1,28}$ = 5.36, $p$ = 0.0282, $d$ = 0.85]. Males, regardless of treatment history, committed significantly more regressive errors compared to never-reinforced errors [Error Type: $F_{2,30}$ = 8.84, $p$ = 0.0010; Never-Reinforced vs. Regressive, $p$ = 0.0142, $d$ = 0.62; Supplemental Figure S3D, significance not shown]. In females, treatment history did not significantly influence trials to reach criterion during any behavioral flexibility task (Supplemental Figure S3A-C). During the set-shift (Supplemental Figure S3D, significance not shown), females committed more perseverative errors relative to never-reinforced errors during the set-shift [$F_{2,29}$ = 8.28, $p$ = 0.0014; Never-Reinforced vs. Perseverative, $p$ = 0.0122, $d$ = 0.22].

**Effort Discounting**

The effects of vapor self-administration during adolescence were also assessed in an effort discounting task in adulthood (Supplemental Figure S4). Both male and female rats showed effort discounting by decreasing their percentage of high reward lever choices as the required lever responses for reinforcement increased [Response Requirement: Males, $F_{4,42} = 339.74$, $p < 0.0001$, range of *d*'s = 0.50 – 1.53; Females, $F_{4,39} = 824.18$, $p < 0.0001$, range of *d*'s = 0.43 – 1.51]; however, treatment did not significantly impact effort discounting for either sex.

**Dendritic Spines in the Medial Prefrontal Cortex**

Unfortunately, after technical issues, only one brain from the male vehicle vapor group was able to be assessed for spine measures. Thus, the sexes were collapsed in the analyses for spine measures (Supplemental Figure S5). Analyses of the proportion of different spine types (Supplemental Figure S5B) revealed no significant effects of treatment but did show a significant effect of spine type [$F_{3,54} = 76.59$, $p < 0.0001$], reflecting that the proportion of the four spine types were all significantly different from each other (*p*'s < 0.05, range of *d*'s = 1.16 – 4.53). Thin spines were the most prevalent, followed by mushroom, stubby, and lastly filopodia. There were no statistically significant effects of treatment on spine density (Supplemental Figure S5C), spine volume, (Supplemental Figure S5D), or spine length (Supplemental Figure S5E).

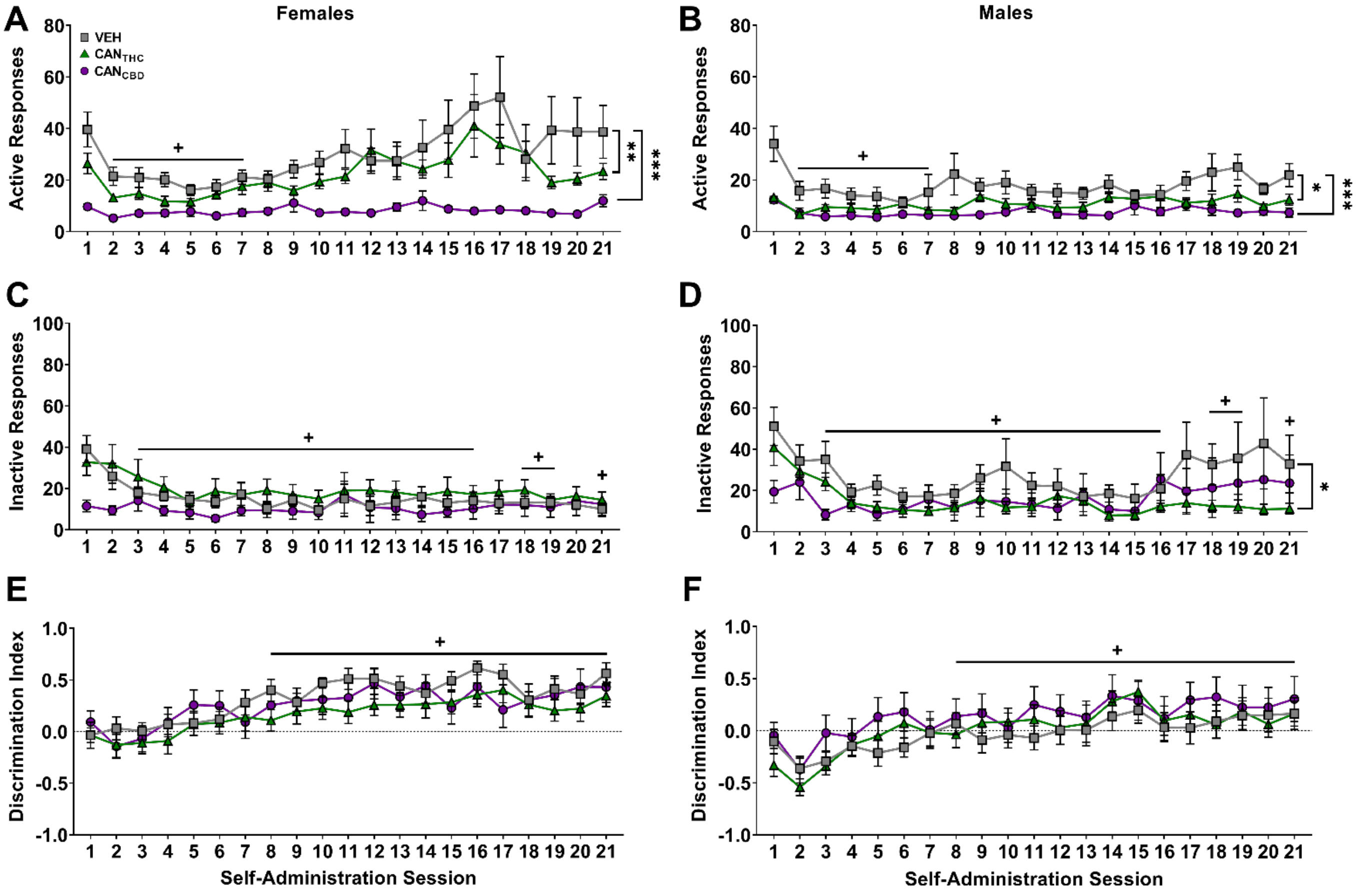


**Supplemental Figure S1**. Nose poke responses in the active port (**A**,**B**), inactive port (**C**,**D**), and nose poke port discrimination indices (**E**,**F**) across 21 vapor self-administration sessions for adolescent female and male rats. Data are shown as mean (± SEM). n = 13-17/group. $^{+}p < 0.05$ vs. session 1, collapsed across treatment and sex; *$p < 0.05$, **$p < 0.01$, ***$p < 0.001$ vs. VEH, collapsed across session.

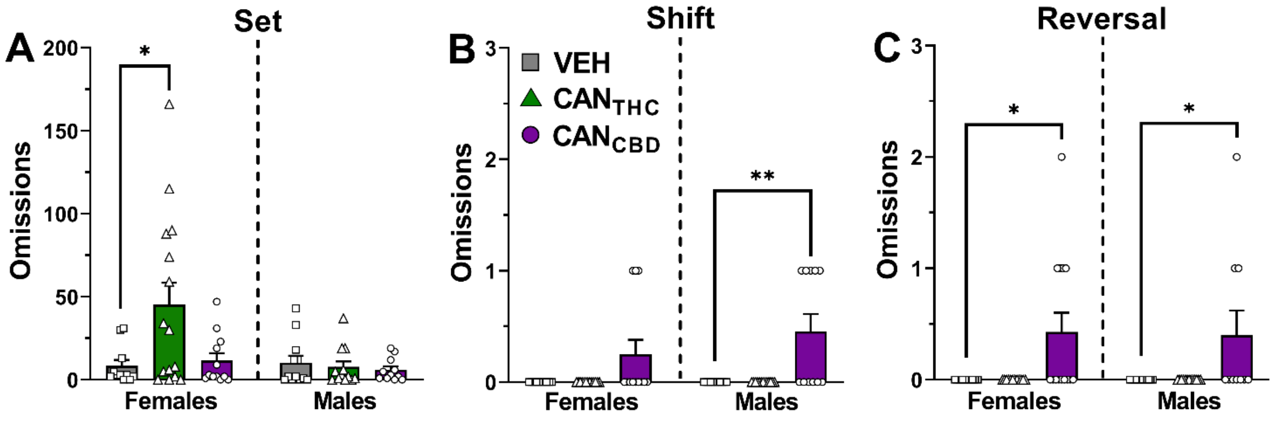


**Supplemental Figure S2**. Omissions committed during different tasks of behavioral flexibility vary depending on sex and treatment. Omissions during the visual cue discrimination (set; **A**), set-shifting (**B**), and reversal (**C**) tasks in the behavioral flexibility test. Data are shown as mean (± SEM). n = 10-15/group. *$p$ < 0.05, **$p$ < 0.01 vs. VEH.

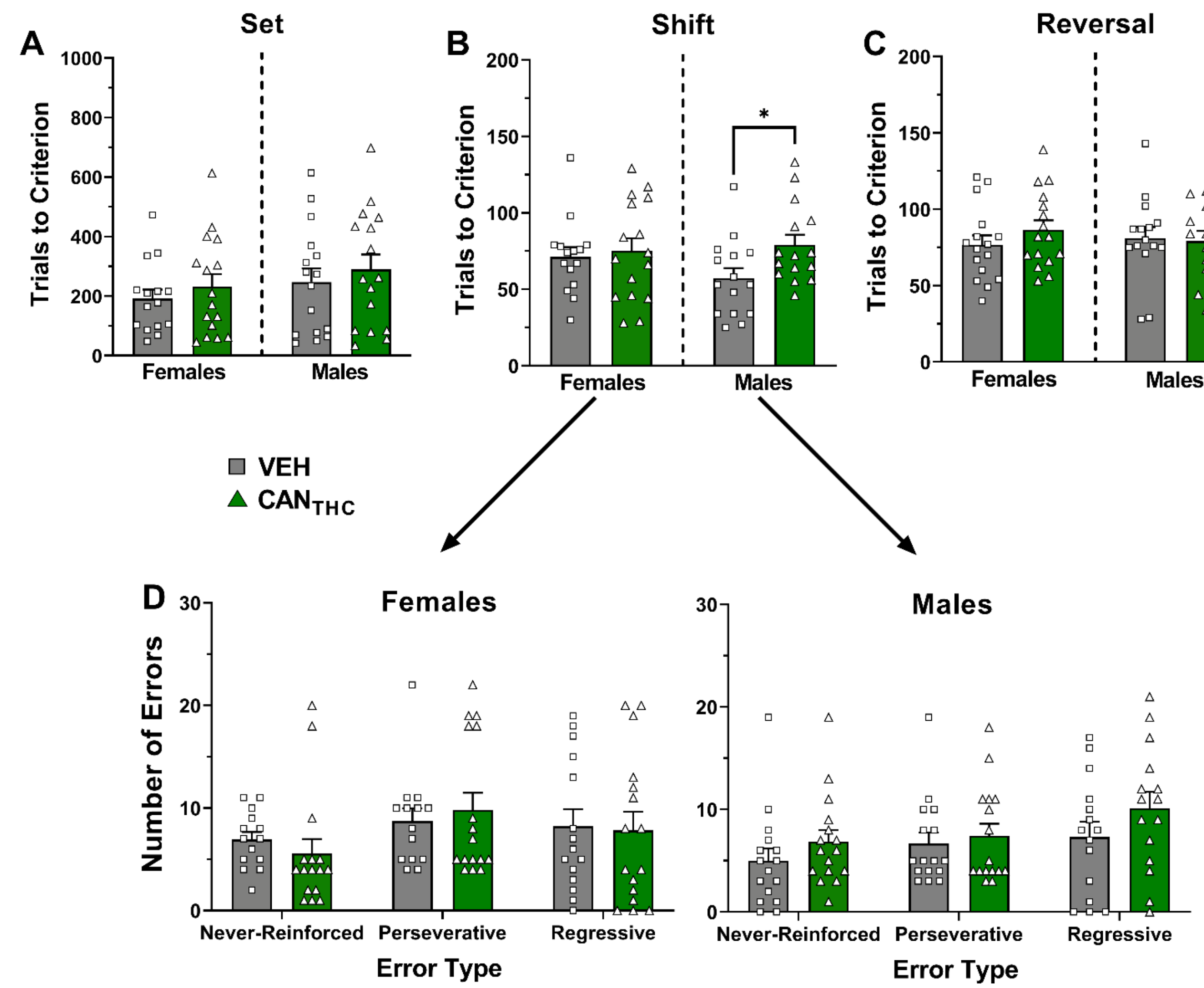


**Supplemental Figure S3**. Passive $CAN_{THC}$ vapor exposure during adolescence impairs behavioral flexibility in male rats when tested in adulthood. Trials to reach criterion in the visual cue discrimination (set; **A**), set-shifting (**B**), and reversal (**C**) tasks in rats of both sexes. (**D**) Number of different types of errors made during the set-shifting task. Data are shown as mean (± SEM). n = 15-16/group. *$p < 0.05$ vs. VEH.

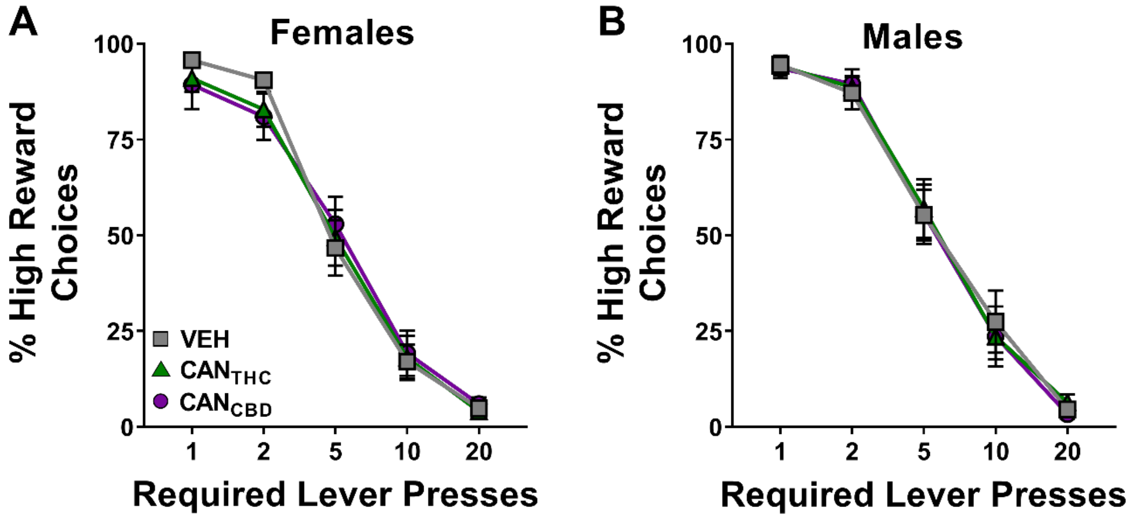


**Supplemental Figure S4**. Vapor self-administration during adolescence does not change effort discounting in adult female (**A**) or male (**B**) rats. Data are shown as mean (± SEM). n = 13-17/group.

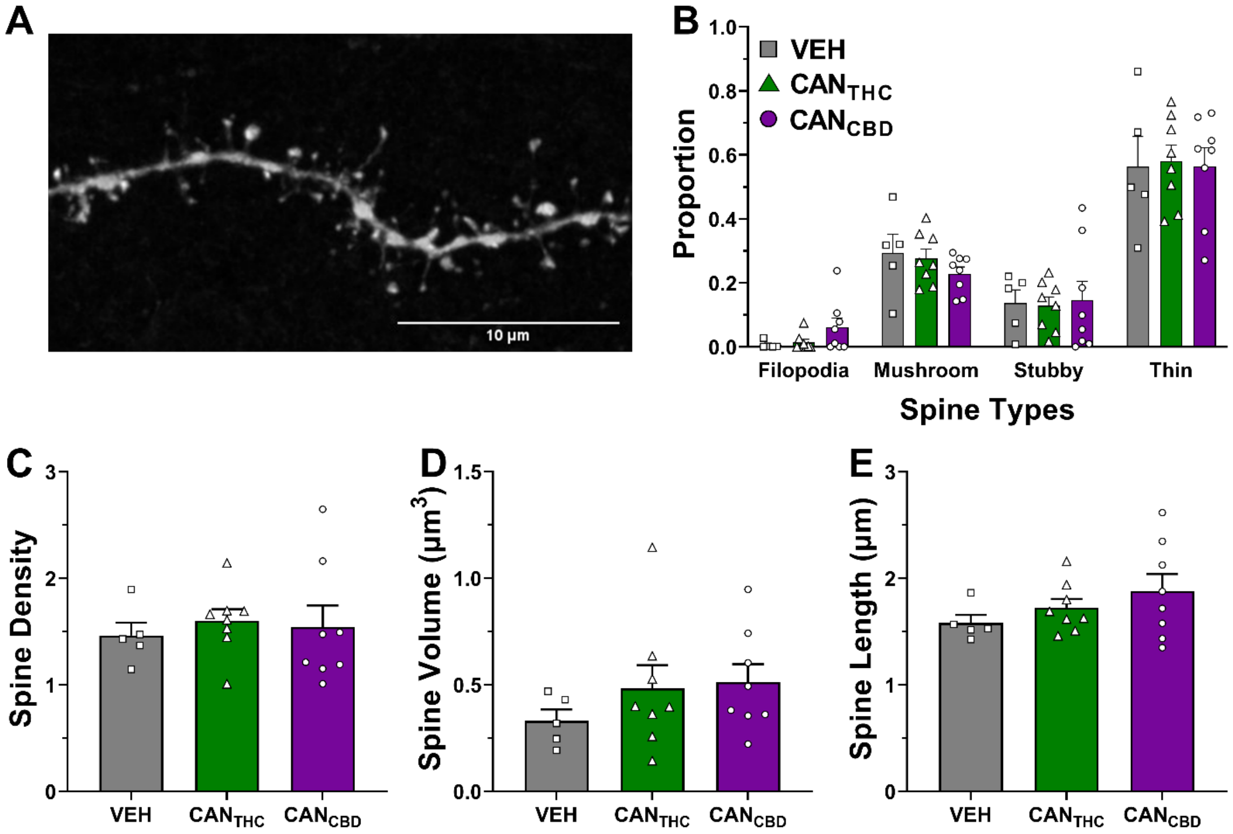


**Supplemental Figure S5**. History of vapor self-administration during adolescence does not significantly impact dendritic spines in the medial prefrontal cortex in adulthood. (**A**) Representative image of a dendrite from the mPFC of a vehicle-exposed female. (**B**) The proportion of the four different types of spines in the mPFC differed significantly regardless of treatment (significance not shown). (**C**) Spine density, (**D**) spine volume, and (**E**) spine length in the mPFC. Data are shown as mean (± SEM). n = 5-8/group (collapsed across sex).
